# Sensory neurons contacting the cerebrospinal fluid require the Reissner fiber to detect spinal curvature *in vivo*

**DOI:** 10.1101/861344

**Authors:** Adeline Orts-Del’Immagine, Yasmine Cantaut-Belarif, Olivier Thouvenin, Julian Roussel, Asha Baskaran, Dominique Langui, Fanny Koëth, Paul Bivas, François-Xavier Lejeune, Pierre-Luc Bardet, Claire Wyart

**Affiliations:** \Institut du Cerveau et de la Moelle épinière (ICM), Sorbonne Université, Inserm U 1127, CNRS UMR 7225, F-75013, Paris, France; Institut Langevin ESPCI, PSL Research University, CNRS UMR7587 1 rue Jussieu, Paris F75005, France

**Keywords:** Cerebrospinal fluid (CSF), CSF-contacting neurons (CSF-cNs), mechanoreception, the Reissner fiber (RF), Polycystin Kidney Disease 2 Like 1 (PKD2L1), SCO-spondin, motile cilia, Kolmer Agduhr cells (KAs), spinal cord, central canal

## Abstract

Recent evidence indicate active roles for the cerebrospinal fluid (CSF) on body axis development and morphogenesis of the spine implying CSF-contacting neurons (CSF-cNs) in the spinal cord. CSF-cNs project a ciliated apical extension into the central canal that is enriched in the channel PKD2L1 and enables the detection of spinal curvature in a directional manner. Dorsolateral CSF-cNs ipsilaterally respond to lateral bending while ventral CSF-cNs respond to longitudinal bending. Historically, the implication of the Reissner fiber (RF), a long extracellular thread in the CSF, to CSF-cN sensory functions has remained a subject of debate. Here, we reveal using electron microscopy in zebrafish larvae that the RF is in close vicinity with cilia and microvilli of ventral and dorsolateral CSF-cNs. We investigate *in vivo* the role of cilia and the Reissner fiber in the mechanosensory functions of CSF-cNs by combining calcium imaging with patch-clamp recordings. We show that disruption of cilia motility affects CSF-cN sensory responses to passive and active curvature of the spinal cord without affecting the Pkd2l1 channel activity. Since ciliary defects alter the formation of the Reissner fiber, we investigated whether the Reissner fiber contributes to CSF-cN mechanosensitivity *in vivo*. Using a hypomorphic mutation in the *scospondin* gene that forbids the aggregation of SCO-spondin into a fiber, we demonstrate *in vivo* that the Reissner fiber *per se* is critical for CSF-cN mechanosensory function. Our study uncovers that neurons contacting the cerebrospinal fluid functionally interact with the Reissner fiber to detect spinal curvature in the vertebrate spinal cord.

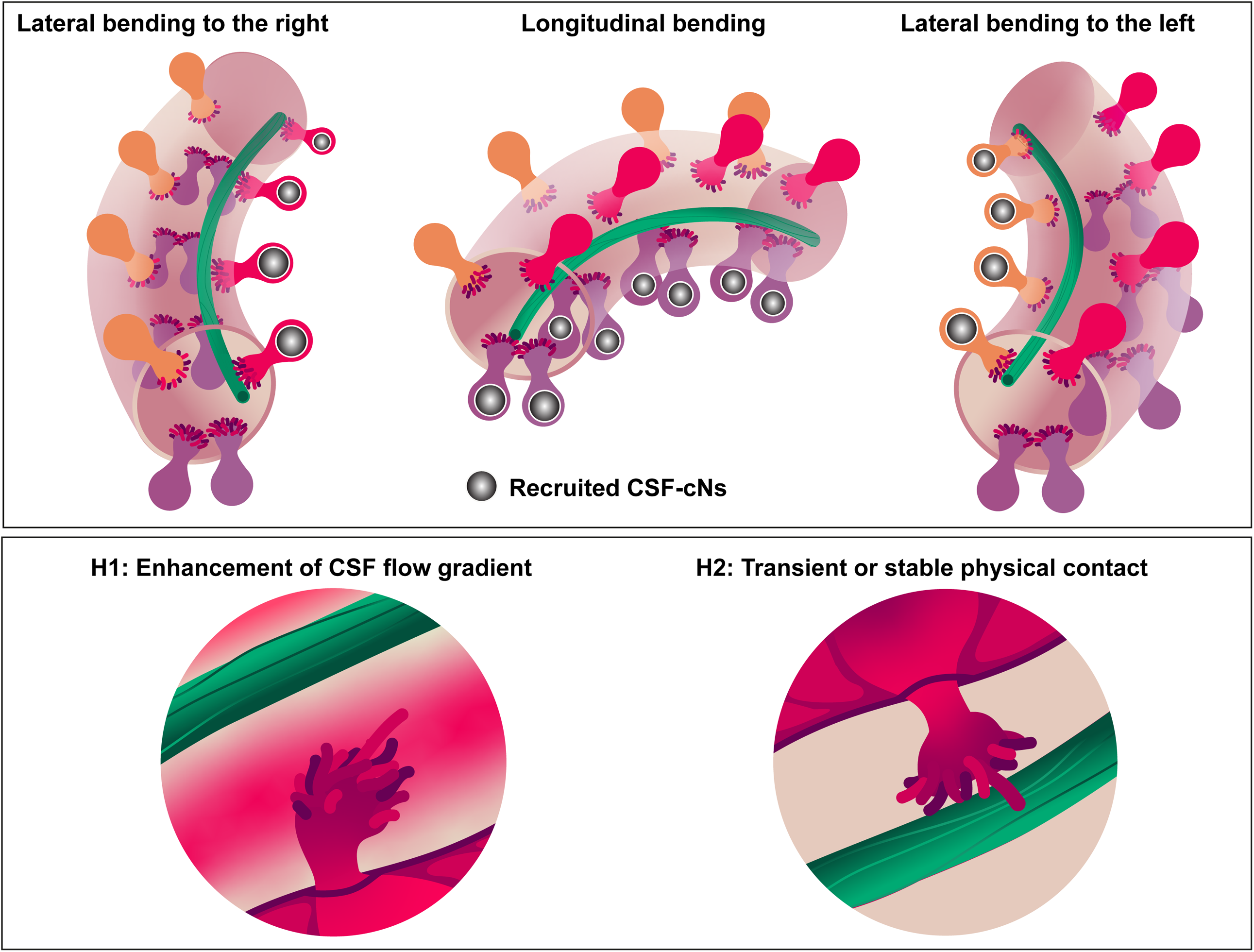

**eToC:** The role of the Reissner fiber, a long extracellular thread running in the cerebrospinal fluid (CSF), has been since its discovery in 1860 a subject of debate. Orts-Del’Immagine *et al*. report that the Reissner fiber plays a critical role in the detection of spinal curvature by sensory neurons contacting the CSF.

**Highlights:**

- Since its discovery, the role of the Reissner fiber has long been a subject of debate
- Mechanoreception in CSF-contacting neurons (CSF-cNs) *in vivo* requires the Reissner fiber
- CSF-cN apical extension is in close vicinity of the Reissner fiber
- CSF-cNs and the Reissner fiber form *in vivo* a sensory organ detecting spinal curvature

## Introduction

Cerebrospinal fluid (CSF) is secreted by the choroid plexuses and fills the ventricular cavities of the brain and the central canal of the spinal cord [1]. The CSF has been long assumed to form a passive fluid acting as a cushion, supporting the clearance of toxic products in the brain, thereby ensuring its mechanical protection and chemical homeostasis. However, multiple studies have shown that secretion and circulation of signaling molecules in the CSF contribute in the brain to neurogenesis in an age-dependent manner [2–7]. Furthermore, physico-chemical properties of the CSF also contribute to organogenesis outside of the nervous system. CSF content and cilia-driven flow control the geometry of the body axis during embryogenesis [8,9] as well as spine curvature in juvenile zebrafish [9,10]. Both the formation of the body axis and the spine organogenesis appear linked to urotensin-related peptides expressed in the spinal cord by cerebrospinal fluid-contacting neurons (CSF-cNs) [9,11].

CSF-cNs are found in the spinal cord in many vertebrate species [12–14]. Spinal CSF-cNs are GABAergic sensory neurons that extend an apical extension into the lumen of the central canal. This apical extension is composed of one motile cilium and numerous microvilli that bath in the CSF [14–20]. Dorsolateral and ventral CSF-cNs originate from two distinct progenitor domains in zebrafish [18,19,21–23] as in mouse [24], and are characterized by different axonal targets [18], and morphology of the apical extension [17]. Both CSF-cN types respond to spinal curvature in a directional manner: while dorsolateral CSF-cNs respond ipsilaterally to lateral bending [16] ventral CSF-cNs are recruited during longitudinal contractions of the spinal cord [25].

Due to the morphological resemblance between CSF-cNs and hair cells, Kolmer had proposed that these cells could constitute a novel sensory organ, referred to as the “SagittalOrgan”, acting as a third ear in the vertebrate spinal cord [13]. This hypothesis has been discussed several times since [12,26–28], but data were based on sparse electron microscopy and not functional evidence. Identifying genetic markers of CSF-cNs over the last decade [29–31] has allowed novel investigation of their function. Based on these recent studies, we know that spinal CSF-cNs sense changes in pH and osmolarity [32–35] as well as mechanical stretch of the spinal cord [16,25,33,36]. Mechanotransduction in CSF-cNs relies on the Polycystic Kidney Disease 2-Like 1 (PKD2L1) channel [16,36], a member of the Transient Receptor Potential (TRP) channel family, which is also a specific marker of these cells in the vertebrate spinal cord [19,30,34,35]. The PKD2L1 channel is enriched in the CSF-cN apical extension [19,36], which differentiates in the larva and likely constitutes the sensory apparatus of CSF-cNs. Concordantly, the length of microvilli in the apical extension tunes the mechanosensory response of CSF-cNs [17].

In the lumen of the central canal, the Reissner fiber (RF) is a long extracellular thread extending caudally from the diencephalic third ventricle to the central canal of the spinal cord and is mainly composed of the aggregation of the SCO-spondin glycoprotein [26,37,38]. A century ago, Kolmer and Agduhr noticed the presence of the RF in close vicinity with the CSF-cN apical extension bathing in the CSF [39]. This proximity led Kolmer to hypothesize a possible functional interaction between the RF and CSF-cNs [13], a hypothesis not favored by Agduhr [12]. Here, we revisited this question by investigating in zebrafish larvae the role of motile cilia and the RF in CSF-cN mechanotransduction.

We show that disruption of cilia motility affects mechanosensory responses of both ventral and dorsolateral CSF-cNs without affecting Pkd2l1 channel activity. We found using electron microscopy that the RF and CSF-cN apical extension come together in close proximity in the lumen of the central canal. We show that in *scospondin* hypomorph lacking the fiber in the central canal, CSF-cNs lose their mechanoreception without disrupting Pkd2l1 channel spontaneous opening. Our results demonstrate the need of the RF for transmitting mechanical deformations associated with spinal curvature to sensory neurons lining the central canal. Altogether, our results validate Kolmer’s hypothesis who suggested that CSF-cNs may functionally interact with the Reissner fiber *in vivo* to form a sensory organ in the vertebrate spinal cord.

## Results

### Motile cilia are required for CSF-cN response to muscle contraction and spinal curvature *in vivo*

We previously showed that cilia functions are necessary for spontaneous activity of CSF-cNs in the embryos [36]. Now, in order to test the role of motile cilia in CSF-cN response to spinal curvature in the larval stage when their apical extension is differentiated, we induced escape responses in 3 days post fertilization (dpf) larvae by puffing artificial cerebrospinal fluid (aCSF) in the otic vesicle (Figure 1). We monitored in the sagittal plane CSF-cN activity with the genetically-encoded calcium sensor GCaMP5G expressed under the CSF-cN specific promoter *pkd2l1* in *Tg(pkd2l1:GCaMP5G)* transgenic larvae [18,29,36] (Figure 1A1, 1A2, 1B2, see also Supplemental Movie S1). In this paradigm, larvae were pinned on the side and muscle contractions subsequent to the otic vesicle stimulation occurred in lateral and horizontal directions, which lead to calcium transients in dorsolateral and ventral CSF-cNs (Figure 1A2 and 1A3). The amplitude of calcium transient in dorsolateral CSF-cNs was larger than in ventral CSF-cNs (mean ΔF / F = 87.2 ± 5 % from 211 dorsolateral CSF-cNs, *vs.* 57.4 ± 3.3 % from 168 ventral CSF-cNs in 15 control sibling larvae; linear mixed model (type II Wald chi-square test): p < 1 × 10^−4^), likely due to the muscle contractions being mainly lateral during the escape. In the *cfap298*^*tm304/tm304*^ mutant larvae with defective motility and polarity of cilia [40–42], the response of CSF-cNs was overall massively reduced (mean ΔF / F = 19.0 ± 1.9 % from 153 dorsolateral CSF-cNs and 15.5 ± 1.4 % from 104 ventral CSF-cNs in 11 mutant larvae; p < 1 × 10^−4^ for dorsolateral CSF-cNs and p < 1 × 10^−3^ for ventral CSF-cNs between mutant and control siblings; Figure 1A2 and 1A3; Supplemental Movie S1 and S2). The proportion of CSF-cNs recruited decreased from 68.2 % to 29.4 % for dorsolateral CSF-cNs and from 64.3 % to 31.7 % for ventral CSF-cNs (Figure 1A3).

**Figure 1.**
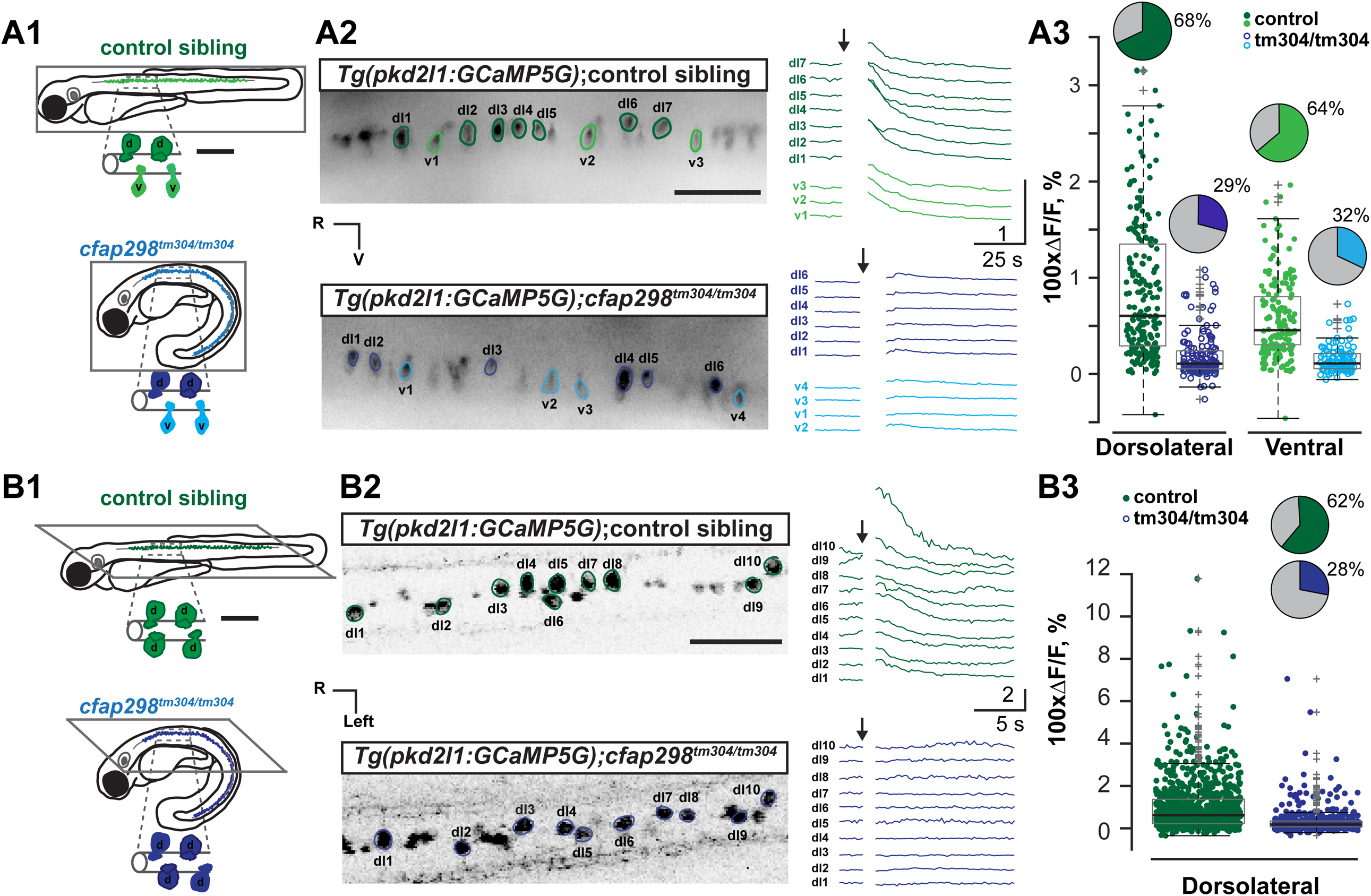
The sensory response of CSF-cNs to spinal curvature is altered in mutants with defective motile cilia. **(A)** CSF-cN calcium transients calculated from the changes in fluorescence of the genetically-encoded calcium indicator GCaMP5G in 3 dpf *Tg*(*pkd2l1:GCaMP5G)* transgenic larvae are large for control siblings (top, green) and strongly reduced in *cfap298* ^*tm304/tm304*^mutants (bottom, blue) during active bends where muscle contraction is induced by pressure-application of aCSF in the otic vesicle (see also **Supplemental Movie S1**; **Supplemental Movie S2**). **(A1)** Diagram representing a 3 dpf larva view in sagittal plane. **(A2)** Time projection stack of 3 optical sections imaged from the sagittal plane (left) showing for illustration purposes a subset of 10 spinal CSF-cNs expressing GCaMP5G in 3 dpf *Tg*(*pkd2l1:GCaMP5G)*. Green and blue lines around soma delimitate ROIs used to calculate ΔF / F traces on the right. **(A3)** Quantification of corresponding ΔF / F in dorsolateral and ventral CSF-cNs in control siblings (green) and *cfap298* ^*tm304/tm304*^ mutants (blue), (mean ΔF/F = 87.2 ± 5.0 % from 211 dorsolateral CSF-cNs and mean ΔF/F = 57.4 ± 3.3 % from 168 ventral cells in 15 control siblings *vs.* mean ΔF / F = 19.0 ± 1.9 % from 153 dorsolateral cells and mean ΔF / F = 15.5 ± 1.4 % from 104 ventral cells in 11 *cfap298* ^*tm304/tm304*^ larvae, linear mixed model (type II Wald chi-square test); p < 1 × 10^−4^ between dorsolateral and ventral CSF-cNs in control siblings, Df = 613; p < 1 × 10^−4^ between dorsolateral CSF-cNs in mutants vs control siblings, Df = 30.4; p < 1 × 10^−3^ between ventral CSF-cNs in mutants vs control siblings, Df = 32.9; p > 0.5 between dorsolateral and ventral CSF-cNs in *cfap*^*298tm304/tm304*^ mutants, Df = 614). Pie charts represent the percentage of responding cells. **(B)** CSF-cN calcium transient recorded in 3dpf *Tg*(*pkd2l1:GCaMP5G)* transgenic larvae after a passive bend of the tail (passive stimulation) in paralyzed control siblings (top, green) and *cfap298*^*tm304/tm304*^ mutant fish (bottom, blue). **(B1)** Diagram showing the coronal plane of a 3 dpf. **(B2)** Time projection stack of 3 optical sections imaged from the coronal plane (left) showing for illustration purposes a subset of 10 dorsolateral CSF-cNs expressing GCaMP5G in 3 dpf *Tg*(*pkd2l1:GCaMP5G)*. Green and blue lines around soma delimitate ROIs used to calculate ΔF / F traces upon passive mechanical stimulation represented on the right. **(B3)** Quantification of corresponding ΔF / F in CSF-cNs represented (as in **(A2)**) during a passive bend (mean ΔF / F = 100.7 ± 4.3 % from 830 cells in 26 control siblings *vs.* 32.3 ± 2.7 % from 490 cells in 15 *cfap298*^*tm304/tm304*^ larvae, linear mixed model (type II Wald chi-square test); p < 2 × 10^−9^; Df = 1; Chi2 = 36.41). Pie charts represent the percentage of responding cells. R, rostral; V, ventral dl: dorsolateral CSF-cNs, v: ventral CSF-cNs. Time projection stacks were constructed from 3 to 4 series of images (corresponding to 0.75 s - 1 s integration time). Each data plotted in **(A3)** and **(B3)** represents one recording from one cell, the central mark on the box plot indicates the median, the bottom and top edges of the box indicate the 25th and 75th percentiles. The whiskers extend to the most extreme data points that are not considered outliers, outliers are identified with a “+” symbol. Scale bars are 500 µm in **(A1)** and **(B1)** and 50 µm in **(A2)** and **(B2)** (left panel).

As the *cfap298* mutation may alter cilia in the otic vesicle, defects in inner ear hair cells could be responsible of the decreased response of CSF-cNs. We therefore examined CSF-cN responses to passive curvature of the spinal cord from paralyzed 3 dpf *Tg(pkd2l1:GCaMP5G)* larvae mounted to record from a coronal view dorsolateral CSF-cNs (Figure 1B1) that are selectively activated by lateral bending of the spinal cord [16,17]. Lateral bending of the tail in control larvae induced as previously reported [16] calcium transients in dorsolateral CSF-cNs (Figure 1B1). In *cfap298*^*tm304/tm304*^ larvae, the response of CSF-cNs was overall massively reduced (mean ΔF / F = 32.30 ± 2.7 % from 490 dorsolateral CSF-cNs in 15 mutant larvae *vs.* 100.7 ± 4.3 % from 830 dorsolateral CSF-cNs in 26 control siblings; p < 2 × 10^−9^; Figure 1B1 and 1B2; Supplemental Movie S2). The proportion of dorsolateral CSF-cNs recruited decreased from 62.2 % to 28.4 % (Figure 1B3). Altogether, our results indicate that cilia motility and polarity are necessary for mechanosensory functions of CSF-cNs *in vivo*.

### Mutants with defective cilia show functional Pkd2l1 channels on CSF-cN apical extension

We previously showed that mechanical activation of CSF-cNs requires Pkd2l1 channels *in vivo* [16] and *in vitro* [36]. To confirm the presence and localization of Pkd2l1 at the apical extension of CSF-cNs in the *cfap298*^*tm304/tm304*^ mutant at 3 dpf, we performed immunohistochemistry in *Tg(pkd2l1:GCaMP5G); cfap298*^*tm304/tm304*^ larvae. Pkd2l1 localized at the apical extension of CSF-cNs in *cfap298*^*tm304/tm304*^ larvae similarly to control (Figure 2A).

**Figure 2.**
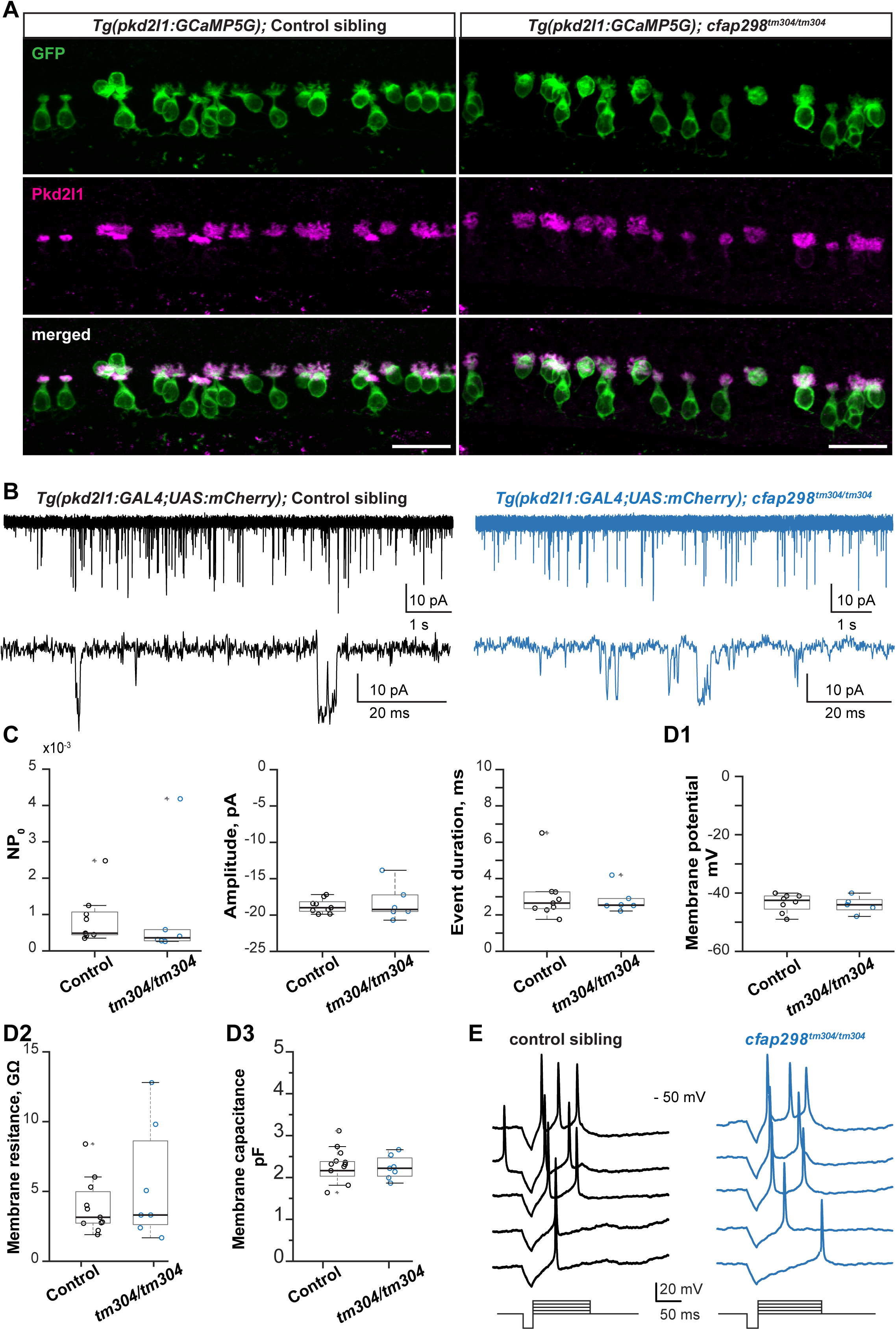
CSF-cNs in *cfap298* mutant with defective motile cilia exhibit functional Pkd2l1 channels. **(A)** Immunohistochemistry with anti-GFP and anti-Pkd2l1 antibodies in 3 dpf *Tg*(*pkd2l1:GCaMP5G)* control sibling (left) and *cfap298*^*tm304/tm304*^ mutant (right) larvae shows in the spinal cord similar localization for Pkd2l1 protein in the apical extension of CSF-cNs (seen in 14 mutant larvae and 8 control siblings). **(B)** *In vivo* whole cell patch-clamp recordings from CSF-cNs using voltage clamp (VC)-mode in 3 dpf *Tg(pkd2l1:GAL4;UAS:mCherry)* show single channel openings in control sibling (left) and in *cfap298*^*tm304/tm3040*^ (right). Bottom traces represent at higher temporal magnification the data from top traces. **(C)** Data plot of unitary current properties show that mutant larvae with defective motile cilia show proper spontaneous channel opening properties (Mean NP_0_ = 9 x10^−4^ ± 2 × 10^−4^ in control siblings *vs.* 1.10^−3^ ± 0.6 × 10^−3^ in mutants, Two-sample Kolmogorov-Smirnov test, p > 0.5, ks2stat = 0.39; Mean unitary current amplitude = - 18.8 ± 0.3 pA for control siblings *vs.* - 18.3 ± 1.0 pA in mutant larvae, p > 0.8, ks2stat = 0.28; Mean duration of single opening = 3.0 ± 0.5 ms in control sibling *vs.* 2.8 ± 0.3 ms in mutant larvae, p > 0.8, ks2stat = 0.28; n = 9 cells in control, n = 6 cells in *cfap298* ^*tm304/tm304*^ larvae). **(D)** Quantification of CSF-cN basic intrinsic electrophysiological properties **(D1)** CSF-cN resting membrane potential is not affected in *cfap298* ^*tm304*^ mutant larvae (Mean membrane potential = - 43.4 ± 1.1 mV in control sibling *vs.* – 44.0 ± 1.3 mV in mutant larvae, Two-sample Kolmogorov-Smirnov test, p > 0.8, ks2stat = 0.30, n = 8 cells in control, n = 5 cells in *cfap298*^*tm304/tm304*^ larvae). **(D2)** Quantification of membrane resistance reveals no change in *cfap298*^*tm304/tm304*^ mutant larvae (Mean membrane resistance = 3.9 ± 0.6 GΩ in control siblings *vs.* 5.5 ± 1.6 GΩ in mutant larvae, Two-sample Kolmogorov-Smirnov test, p > 0.8, ks2stat = 0.29; n = 11 cells in controls, n = 7 cells in *cfap298*^*tm304/tm304*^ larvae) **(D3)** CSF-cN membrane capacitance is not affected in *cfap298*^*tm304*^ mutant larvae (Mean membrane capacitance = 2.2 ± 0.1 pF in control sibling *vs.* 2.1 ± 0.1 pF in mutant larvae, Two-sample Kolmogorov-Smirnov test, p > 0.2; ks2stat = 0.44, n = 11 cells in control, n = 7 cells in *cfap298*^*tm304/tm304*^ larvae). **(E)** CSF-cN action potential discharge recorded in current clamp (CC)-mode in response to successive current steps (100 ms-long pulses from 2 pA to 10 pA in 2 pA increments). NP_0_, opening probability. Each data point represents one recording from one cell; plots use median as measure of central tendency (central mark on the box plot), bottom and top edges of the box indicate the 25th and 75th percentiles. The whiskers extend to the most extreme data points without considering outliers, which are identified with a “+” symbol. Scale bars is 20 µm in **(A)**.

In order to confirm the functionality of Pkd2l1 channels in CSF-cNs, we performed *in vivo* whole-cell voltage clamp recordings in 3 dpf *Tg(pkd2l1:GAL4;UAS:mCherry); cfap298*^*tm304/tm304*^ larvae. We observed spontaneous Pkd2l1 channel openings with similar properties in both *cfap298*^*tm304/tm304*^ mutants and their control siblings (Figure 2B, 2C Two-sample Kolmogorov-Smirnov test: p > 0.5). Furthermore, we did not notice any effect of cilia defects on CSF-cN passive properties (Figure 2D1, 2D2, 2D3, p > 0.2) or firing patterns upon current injection (Figure 2E). Hence, cilia impairment decreases CSF-cN mechanosensory function without affecting their intrinsic excitability nor Pkd2l1 spontaneous channel openings.

### Mechanosensory function of CSF-cNs requires the Reissner fiber

The c*fap298*^*tm304*^ mutation affecting cilia polarity and motility [40,42] has been shown to *i)* reduce CSF flow [10,43] and CSF transport [36], *ii)* reduce the diameter of the central canal [43] and *iii)* forbids the formation of the RF [8]. Parameters such as CSF flow, diameter of the central canal and the presence of the RF could all contribute to the defect observed in CSF-cN sensory function.

Since the role of the RF in spinal mechanoreception is of peculiar interest, we took advantage of the hypomorphic mutation *scospondin*^*icm15*^ in the gene *scospondin* in which a 5 amino acid insertion in the EMI domain forbids the aggregation of SCO-spondin into the Reissner fiber [8]. Response of dorsolateral and ventral CSF-cN to active tail bending was largely abolished in *Tg(pkd2l1:GCaMP5G); scospondin*^*icm15/icm15*^ mutant larvae (Figure 3A1, A2, Supplemental Movie S3; dorsolateral CSF-cNs: mean ΔF / F = 57.3 % ± 3.0 % from180 cells in 12 control sibling, *vs*. 7.1 % ± 0.6 % from 167 cells in 11 mutant larvae, linear mixed model (type II Wald chi-square test): p < 1 × 10^−4^; ventral CSF-cNs: mean ΔF / F = 31.3 ± 2.1 % from 128 cells in control sibling, *vs*. 10.9 ± 1.4 % from 146 cells from mutant larvae, linear mixed model (type II Wald chi-square test): p < 5 × 10^−4^). In response to passive tail bending (Figure 3B1), the calcium transients recorded in dorsolateral CSF-cNs showed a threefold reduction in *scospondin*^*icm15/icm15*^ mutant larvae (Figure 3B2, mean ΔF / F = 194.0 ± 17.8 % from 276 cells in 11 control siblings, *vs*. 56.2 ± 13.1 % from 165 cells from in mutant larvae, p < 0.005). In the active assay, the proportion of dorsolateral CSF-cNs recruited was decreased by 90.8 % and by 77.8 % for ventral CSF-cNs (Figure 3A2). After passive stimulation, the proportion of responding dorsolateral CSF-cNs was decreased by 72.1 %. (Figure 3B2). Altogether, our results demonstrate that CSF-cNs require the RF to optimally respond to mechanical stimuli associated with spinal curvature *in vivo*.

**Figure 3.**
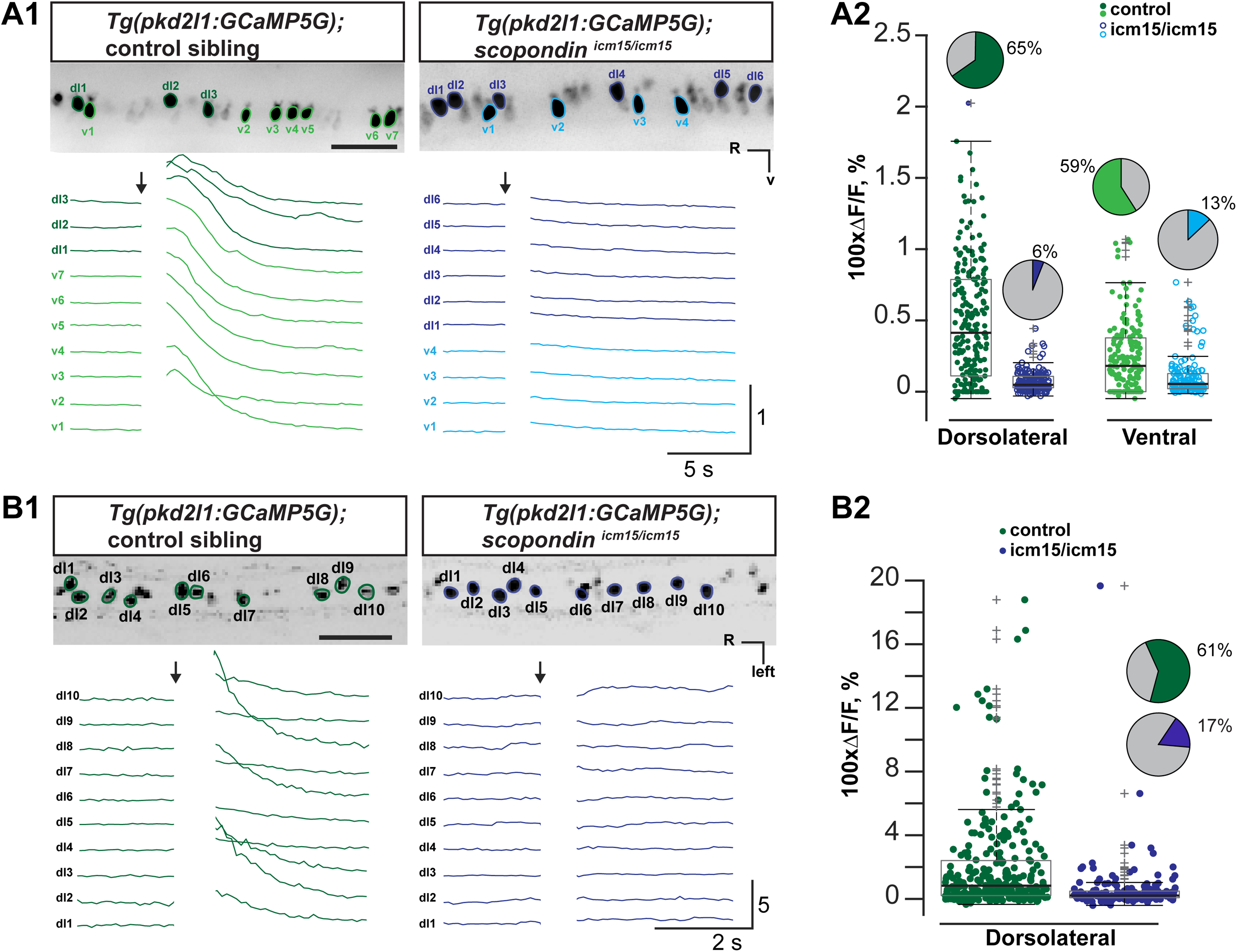
The absence of the Reissner fiber reduces the sensory response of CSF-cNs to spinal curvature. **(A)** CSF-cN calcium transient recorded in 3dpf *Tg*(*pkd2l1:GCaMP5G)* after active bends due to muscle contraction induced by pressure-application of aCSF in the otic vesicle in *scospondin*^*icm15/icm15*^ mutant larvae (right, blue) and control sibling (left, green) (see also **Supplemental Movie S3**). **(A1)** Time projection stack of 3 optical sections imaged from the sagittal plane showing dorsolateral and ventral CSF-cNs expressing GCaMP5 (top) in the spinal cord with ROIs on somas (green and blue line) used to calculate ΔF / F traces below. **(A2)** Quantification of calcium transients in dorsolateral and ventral CSF-cNs in control and *scospondin*^*icm15/icm15*^ mutants in control sibling (green circle) and mutant larvae (blue circle), (mean ΔF / F = 57.3 ± 3.0 % from 180 dorsolateral cells and mean ΔF / F = 31.3 ± 2.1 from 128 ventral cells in 12 control siblings *vs.* mean ΔF / F = 7.1 ± 0.6 % from 167 dorsolateral cells and mean ΔF / F = 10.9 ± 1.4 % from 146 ventral CSF-cNs in 11 *scospondin*^*icm15/icm15*^ larvae, linear mixed model (type II Wald chi-square test); p < 1 × 10^−4^ between dorsolateral and ventral CSF-cNs in control sibling; Df = 602; p < 1 × 10^−4^ between dorsolateral CSF-cNs in control siblings *vs.* mutant larvae; Df 21.4; p < 5 × 10^−4^ between ventral CSF-cNs in control siblings *vs.* mutant larvae; Df 22.6; p>0.05 between dorsolateral and ventral CSF-cNs in *scospondin*^*icm15/icm15*^ larvae, Df 601). Pie charts represent the percentage of responding cells. **(B)** CSF-cN calcium transient after a passive bend of the tail in paralyzed control siblings (left, green) and *scospondin* ^*icm15/icm15*^ mutant (right, blue) larvae. **(B1)** Time projection stack of 3 optical sections imaged from the coronal plane shows CSF-cNs expressing GCaMP5 protein (top). ROIs (green and blue lines) used to calculate ΔF / F traces upon passive mechanical stimulation. **(B2)** Same as **(A2)** during passive bending (mean ΔF / F = 194.0 ± 17.8 % from 276 cells in 11 control siblings *vs.* mean Δ F / F = 56.2 ± 13.1 % from 165 cells in 8 *scospondin*^*icm15/icm15*^ larvae, linear mixed model (type II Wald chi-square test); p < 5 x 10^−3^; Df = 1; Chi2 = 8.18). Scale bars are 50 µm in **(A1)** and **(B1)**. dl: dorsolateral CSF-cNs, R, rostral; V, ventral, v: ventral CSF-cNs. Z-projection stacks were constructed from 3 series of images (corresponding to 0.75 s - 1 s integration time). For box plots, each data point in **(A2)** and **(B2)** represents one recording from one cell, the central mark indicates the median, the box bottom and top edges indicate the 25th and 75th percentiles.

### Pkd2l1 channels in CSF-cNs remain functional when the Reissner fiber is absent

The RF could be required for CSF-cNs to express the Pkd2l1 channel at the membrane. To verify that CSF-cNs properly express Pkd2l1 in their apical extension, we first performed immunohistochemistry on *Tg(pkd2l1:GCaMP5G); scospondin*^*icm15/icm15*^ and found that the Pkd2l1 protein was still enriched in CSF-cN apical extension of larvae lacking the RF (Figure 4A). Accordingly, whole-cell patch-clamp recording of CSF-cNs in *Tg(pkd2l1:GAL4;UAS:mCherry); scospondin*^*icm15/icm15*^ revealed spontaneous unitary current (Figure 4B-C, Two-sample Kolmogorov-Smirnov test: p > 0.8), most likely reflecting functional Pkd2l1 channels [34,36]. We tested whether the absence of the RF could affect the intrinsic properties of CSF-cNs and found no difference in membrane resistance, membrane capacitance, or resting membrane potential (Figure 4D1-3; p > 0.07). The firing pattern of CSF-cNs in *scospondin*^*icm15/icm15*^ upon current injection was comparable to controls (Figure 4E). Altogether, our results show that the absence of the RF does not alter the localization of Pkd2l1 channel to the apical extension, the spontaneous Pkd2l1 channel properties nor the intrinsic excitability of CSF-cNs.

**Figure 4.**
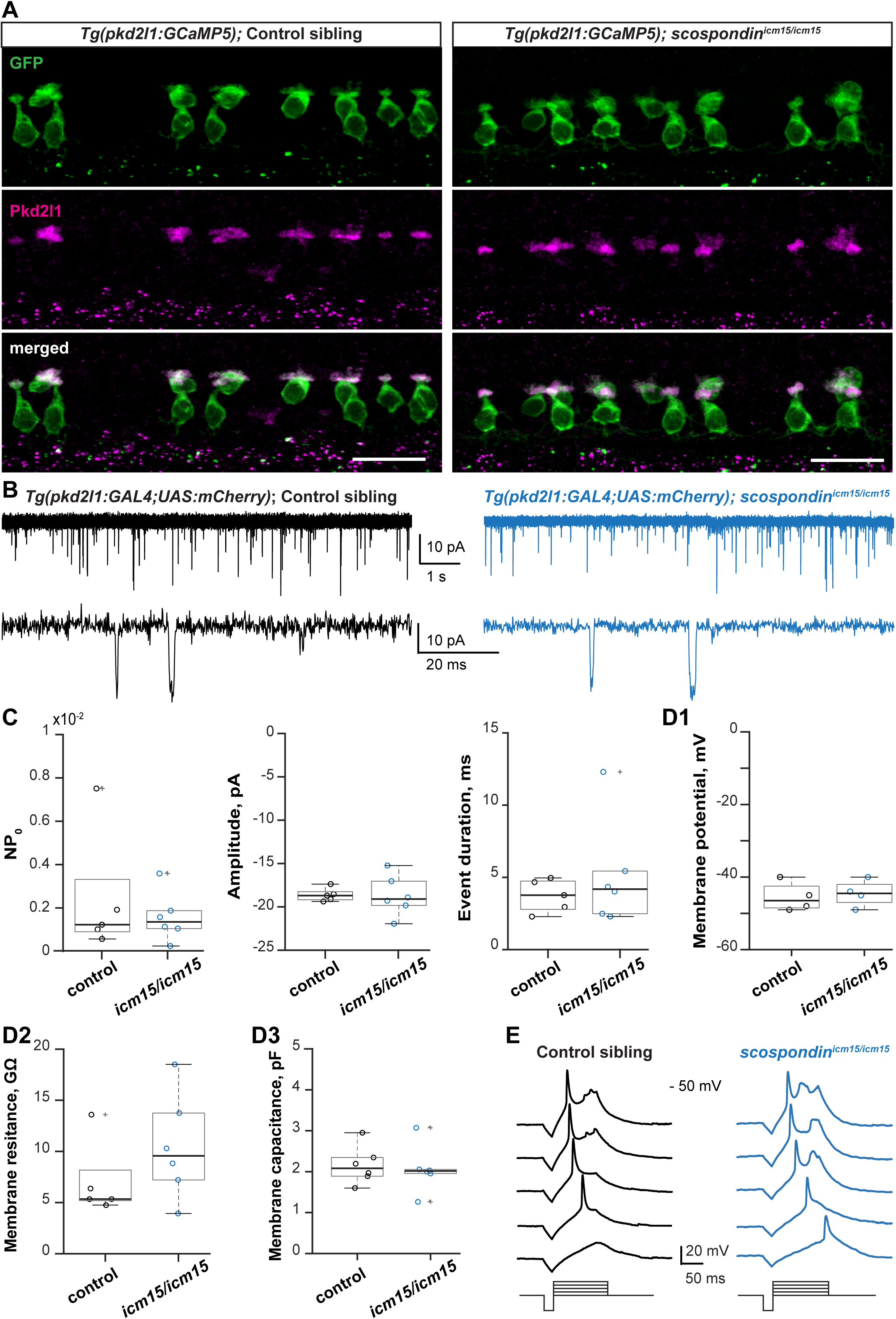
*scospondin* mutants deprived of the Reissner fiber exhibit functional Pkd2l1 channels. **(A)** Immunohistochemistry for GFP and Pkd2l1 in 3dpf *Tg*(*pkd2l1:GCaMP5G)* control siblings (left) and *scospondin*^*icm15/icm15*^ mutant larvae (right) shows that Pkd2l1 protein is localized in CSF-cN apical extension (sagittal view, n = 15 control sibling and 9 mutant larvae). **(B)** *In vivo* whole cell patch-clamp recording of CSF-cNs in voltage clamp (VC)-mode targeted for mCherry in 3dpf *Tg(pkd2l1:GAL4;UAS:mCherry)* exhibits single channel opening in control sibling (left) and in *scospondin*^*icm15/icm15*^ mutant (right) larva. Bottom traces represent a higher magnification from the top trace. **(C)** Unitary currents reflecting spontaneous channel opening properties are not affected in *scospondin*^*icm15/icm15*^ mutants compared to control siblings (Mean NP_0_ = 2.4 × 10^−3^ ± 1.3 × 10^−3^ in control sibling *vs.* 1.6 × 10^−3^ ± 0.5 × 10^−3^ in mutant, Two-sample Kolmogorov-Smirnov test, p > 0.9, ks2stat = 0.23; Mean unitary current amplitude = - 18.6 ± 0.3 pA in control sibling *vs.* −18.7 ± 0.9 pA in mutant larvae, p > 0.8, ks2stat = 0.33; Mean duration of single opening = 3.7 ± 0.5 ms in control sibling *vs.* 5.1 ± 1.5 ms in mutant larvae, p > 0.8, ks2stat = 0.33; n = 5 cells in control siblings, n = 6 cells in *scospondin*^*icm15/icm15*^ larvae). **(D)** Quantification of CSF-cN basic intrinsic electrophysiological properties **(D1)** CSF-cN resting membrane potential remains unaffected in *scospondin*^*icm15/icm15*^ mutants (Mean membrane potential = - 45.5 ± 2.0 mV n = 4 cells in control sibling *vs.* – 44.5 ± 1.8 mV, n = 4 cells in mutant larvae, Two-sample Kolmogorov-Smirnov test, p > 0.9, ks2stat = 0.25). **(D2)** CSF-cN membrane resistance is not altered in *scospondin*^*icm15/icm15*^ mutants (Mean membrane resistance = 6.9 ± 1.4 GΩ in control sibling *vs.* 10.4 ± 2.1 GΩ in mutant larvae, Two-sample Kolmogorov-Smirnov test, p > 0.08, ks2stat = 0.67; n = 6 cells in control, n = 6 cells in *scospondin*^*icm15/icm15*^ larvae). **(D3)** CSF-cN membrane capacitance is comparable between *scospondin*^*icm15/icm15*^ mutants and control larvae (Mean membrane capacitance = 2.2 ± 0.2 pF n = 6 cells in control sibling *vs.* 2.1 ± 0.2 pF n = 6 cells in *scospondin*^*icm15/icm15*^ larvae, Two-sample Kolmogorov-Smirnov test, p > 0.8, ks2stat = 0.33). **(E)** Discharge of action potential in CSF-cNs recorded in current clamp (CC)-mode in response to successive current steps (100 ms long-pulses from 2 pA to 10 pA in 2 pA increments). Sale bars is 20 µm in **(A)**. NP_0_, opening probability. Each data point represents one recording from one cell; plots use median as measure of central tendency (central mark on the box plot), the bottom and top edges of the box indicate the 25th and 75th percentiles. The whiskers extend to the most extreme data points without considering outliers, and the outlier is identified with a “+” symbol.

### In the central canal, the Reissner fiber is in close vicinity with CSF-cN apical extension

Given that in the absence of the RF, CSF-cN sensitivity to spinal curvature is largely reduced despite the cells retaining their intrinsic properties and Pkd2l1 channel activity, we investigated where the RF is localized in the central canal relative to the apical extension of CSF-cNs. In live larvae, the lumen of the central canal is typically 8.7 ± 0.4 µm width and 10.2 ± 0.7 µm height (measured from 4 and 9 larvae respectively, Figure 5A, 5C and 5D) and CSF-cNs extend their dendritic apical extension by typically 2.9 ± 0.1 µm height (measured from 9 cells in 4 larvae) towards the center of the central canal, suggesting that CSF-cNs cover a substantial portion of the central canal in living zebrafish larvae.

**Figure 5.**
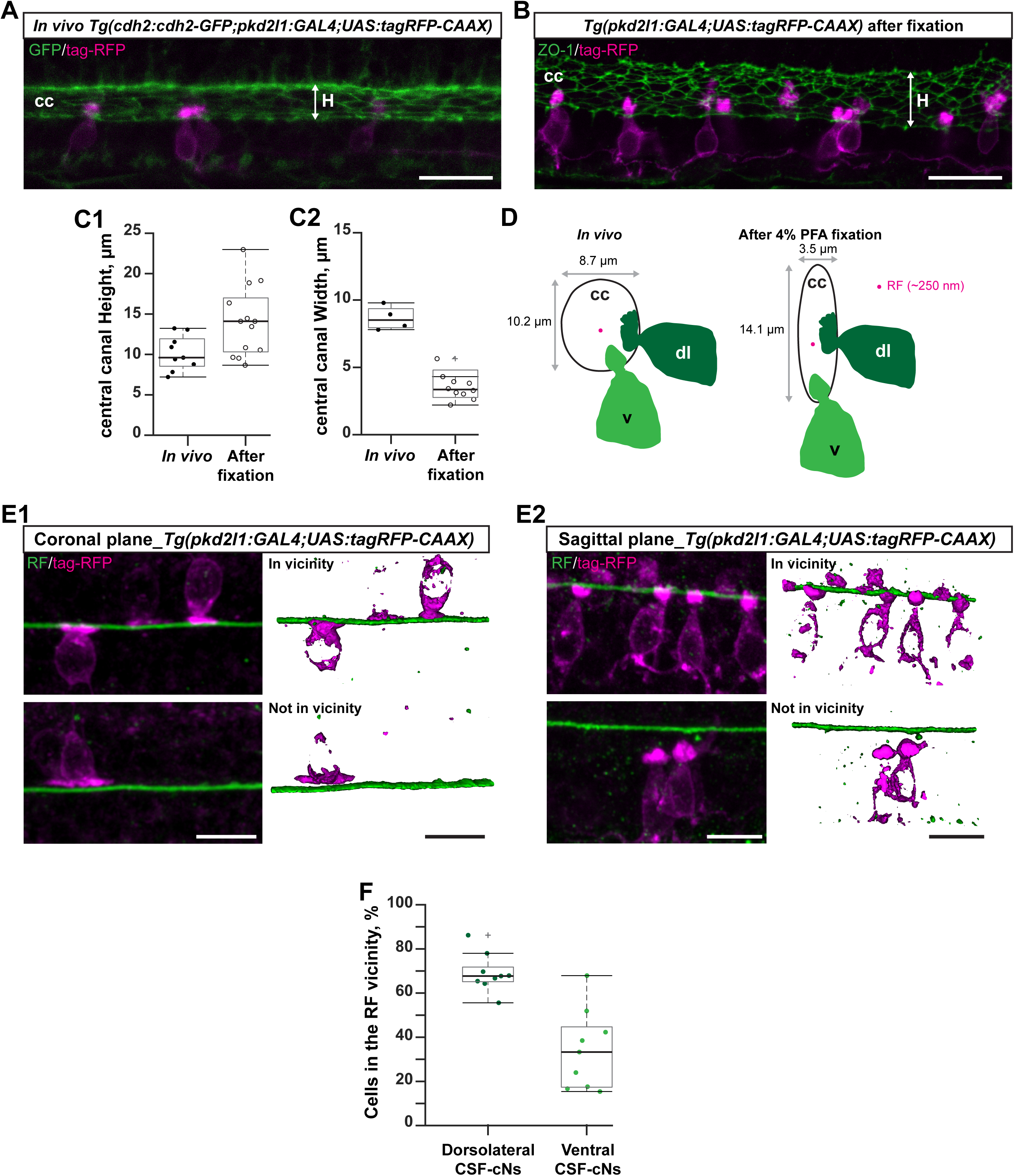
The Reissner fiber is in close vicinity of the apical extension of CSF-cNs. **(A)** Z projection stack of a few optical sections imaged in the sagittal plane in living 3 dpf *Tg(cdh2:cdh2-GFP; pkd2l1:GAL4;UAS:tagRFP-CAAX)* larvae shows the height of the central canal *in vivo*. **(B)** Z projection stack of a few optical sections imaged in the sagittal plane in 3 dpf *Tg(pkd2l1:GAL4;UAS:tagRFP-CAAX)* larvae after co-immunostaining for ZO-1 and tag-RFP used to measure the height of the central canal in fixed whole mount larva. **(C)** Comparison of the size of the central canal between live and fixed whole mount 3 dpf larva reveals an artifact of fixation. **(C1)** Measurement of the height of the central canal shows that the central canal has a higher height after fixation (mean height = 10.2 ± 0.7 µm from 9 live larvae *vs.* 14.1 ± 1.2 µm in 13 fixed whole mount larvae, Two-sample Kolmogorov-Smirnov test, p < 0.03, ks2stat = 0.62). **(C2)** Measurement of the width of the central canal reveals that the central canal is two times narrower after fixation (mean width = 8.7 ± 0.4 µm from 4 live larvae *vs*. 3.5 ± 0.3 µm in 10 fixed whole mount larvae, Two-sample Kolmogorov-Smirnov test, p < 2 x 10^−3^, ks2stat = 1). **(D)** Diagram representing the effect of 4 % PFA fixation on the shape of the central canal. **(E)** Z projection stack of a few optical sections imaged after immunohistochemistry for tag-RFP and Reissner material in 3 dpf *Tg(pkd2l1:GAL4;UAS:tagRFP-CAAX)* larvae reveals that the apical extension of dorsolateral and ventral CSF-cNs are in close vicinity of the RF. **(E1)** Dorsolateral CSF-cNs observed in coronal plane are found in the vicinity of the RF (top panel, In vicinity) or further away (bottom panel, Not in vicinity). Panel on the right represents a 3D reconstruction from the Z-stack on the left. **(E2)** Same as **(E1)** imaged in the sagittal plane in order to visualize the proximity of ventral CSF-cNs with the RF. **(F)** Dorsolateral CSF-cNs imaged in the coronal plane more often appear in close proximity to the RF than ventral CSF-cNs imaged in sagittal plane (mean fraction in vicinity = 69.0 ± 2.9 % from 281 dorsolateral CSF-cNs in 9 larvae *vs.* 34.2 ± 6.0 % from 213 ventral CSF-cNs in 9 larvae; Two-sample Kolmogorov-Smirnov test, p < 5 × 10^−4^; ks2stat = 0.89). Sale bar is 20 µm in **(A)** and **(B)** and 10 µm in **(E)** and **(F)**. cc: central canal; dl: dorsolateral CSF-cNs; H: Height; RF: the Reissner fiber; v: ventral CSF-cNs. Each data point represents one measure from one fish; plots use median as measure of central tendency (central mark on the box plot), the bottom and top edges of the box indicate the 25th and 75th percentiles. The whiskers extend to the most extreme data points without considering outliers, and the outlier is identified with a “+” symbol.

To assess the relative organization of the RF and CSF-cN apical extension, we immuno-stained for the Reissner fiber material as previously described [8] in 3 dpf *Tg(pkd2l1:GAL4;UAS:tagRFP-CAAX)* larvae. As the process of fixation can alter the shape of cavities filled with CSF such as the central canal, we determined the impact of fixation on the width and height of the central canal measured in 3 dpf *Tg(cdh2:cdh2-GFP; pkd2l1:GAL4;UAS:tagRFP)* larvae imaged live and after fixation followed by either co-immunostaining for GFP and tag-RFP, or ZO-1 and tag-RFP (Figure 5A-D). PFA fixation induced a shrinking of the central canal along the midline (Figure 5A and 5B): in the transverse plane, the lumen of the central canal after fixation became narrower (3.5 ± 0.3 µm, Two-sample Kolmogorov-Smirnov test: p < 0.05) and more elongated along the dorsoventral axis (14.1 ± 1.2 µm, p < 0.002) (Figure 5C and 5D). In these conditions, we investigated the approximate position of the RF relative to the apical extension of dorsolateral (Figure 5E1) and ventral (Figure 5E2) CSF-cNs in 3 dpf *Tg(pkd2l1:GAL4;UAS:tagRFP-CAAX)* larvae co-immunostained for RF and tag-RFP. The RF was in close vicinity of two third of dorsolateral CSF-cNs (Figure 5E1, top panel and 5F) and one third of ventral CSF-cNs (Figure 5E2, top panel and 5F; Two-sample Kolmogorov-Smirnov test: p < 5 × 10^−4^).

At higher resolution, we characterized the ultrastructure of the central canal in sagittal sections of 3-4 dpf wild type larvae using transmission electron microscopy (Figure 6). In single sections (Figure 6A, 6B) and reconstruction of the RF and cilia in 3D using serial block face scanning electron microscope imaging (Figure 6C, 6D), the RF appeared as a long and thin thread (Figure 6A, 5B) with diameter of 258.4 nm ± 6.8 nm (26 measurements from 2 larvae) often in close contact with cilia (arrows) and microvilli (arrowheads) (Figure 6A). Cilia in contact with the RF had two central microtubules along the axoneme (Figure 6A1, 6A2, 6A3, 6A4), which typically characterize motile cilia found in ependymal radial glia [36,44–46] as well as CSF-cNs [16]. CSF-cNs have one motile cilium and many microvilli [16–18]. In these ultrastructure images, a subset of dorsolateral and ventral CSF-cNs extended toward the RF via both their microvilli (Figure 6A1, 6B, arrowheads) and their motile cilium (Figure 6B-D, see also Supplemental Movie S4).

**Figure 6.**
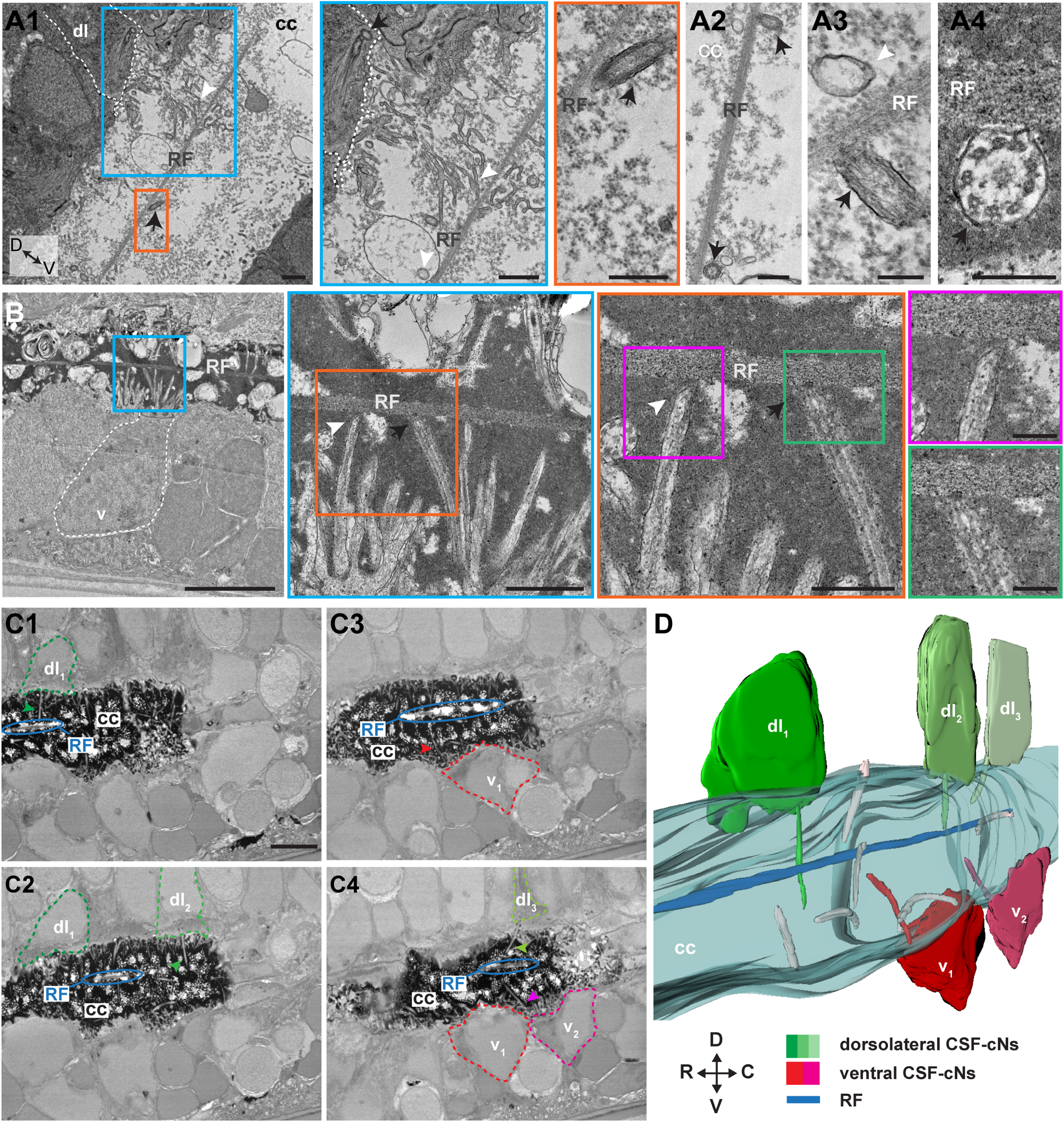
The Reissner fiber can contact cilia and microvilli of CSF-cNs in the central canal. **(A, B)** Transmission electron microscopy (TEM) from spinal cord sectioned in the sagittal plane reveals motile cilium (black arrow) near or in contact with the Reissner fiber (RF) as well as microvilli from CSF-cNs (white arrowhead) near RF. Zoomed regions highlighted in different colors. **(A1)** The RF tends to be dorsally-located in the central canal and close to cilia (arrow), as well as microvilli (arrowhead) originating from identifiable dorsolateral CSF-cNs. **(A2), (A3), (A4)** Appositions onto the RF of motile cilia doted of two central microtubules along the axoneme (arrow in **(A2), (A4)**) as well as microvilli from a ventral CSF-cN (arrowhead, **(A3)**). **(B)** Zooms of lateral regions delineated in colored lines show that ventral CSF-cNs can apparently contact the RF via their microvilli (arrowheads), and their motile cilium (arrows). **(C)** Z projection stack of 3 subsequent images acquired in the sagittal plane from serial block face scanning electron microscopy (SBF-SEM) highlighting apparent contacts between RF (blue ellipse) and dorsolateral CSF-cNs. **(C1), (C2), (C3), (C4)** correspond to different position in the sagittal plane where multiple dorsolateral and ventral CSF-cNs can be observed along the rostro-caudal axis (see **Supplemental Movie S4**). **(D)** 3D reconstruction after SBF-SEM imaging (60 sections of 40 nm Z steps and 7 nm thickness) shown in **(C)** and in **Supplemental Movie S4.** Scale bars are 1 µm in **(A1)** and **(A2)**, 200 nm in **(A3)** and **(A4)**, 2 µm in **(B)**, 1 µm in blue line delineated in **(B)**, 500 nm in orange line delineated in **(B)**, 200 nm in pink and green lines delineated in **(B)** and 5µm in **(C)**. C: caudal, cc: central canal, D: dorsal, dl: dorsolateral CSF-cNs, V: ventral: v: ventral CSF-cNs, R: rostral, RF: Reissner’s fiber. Note that the background signal in the central canal obtained by TEM varies depending on the fixation methods: light background signal for PFA / TCA in **(A1), (A2)** and **(A3)** and dense background signal for PFA/Glutaraldehyde in **(A4), (B)** and **(C)**.

## Discussion

Our study demonstrates *in vivo* the functional coupling between the Reissner fiber and neurons contacting the cerebrospinal fluid in the spinal cord in order to detect spinal curvature. By analyzing the physiological and sensory properties of CSF-cNs in the *scospondin*^*icm15*^ mutant lacking the Reissner fiber, we proved that Kolmer was correct when he speculated that neurons contacting the cerebrospinal fluid form a mechanosensory organ with the Reissner fiber in the vertebrate spinal cord [13].

### The Reissner fiber functionally interacts with CSF-cNs to sense spinal curvature *in vivo*

We had formerly showed that CSF-cNs integrate *in vivo* mechanosensory inputs on the concave side during spinal curvature [16,17] in order to respond to lateral bending for dorsolateral CSF-cNs [16,17] and longitudinal bending for ventral CSF-cNs [25]. *In vivo*, this directional mechanosensory response of CSF-cNs requires the channel Pkd2l1 [16]. Recently, we showed that CSF-cNs when isolated *in vitro* keep their mechanosensory properties: the open probability of the channel Pkd2l1 is largely modulated by mechanical pressure on CSF-cN membrane [36]. Now, we made a new step in understanding how CSF-cNs detect curvature *in vivo*: we demonstrate that the RF enhance by at least threefold the response of CSF-cNs to spinal curvature *in vivo*.

Note that in *cfap298* mutants, the disruption of cilia motility leads to the absence of the Reissner fiber [8] together with a reduction in the dimensions of the central canal lumen [43]. In these mutants, we cannot exclude that the narrow central canal lumen could also contribute with the loss of Reissner fiber to the reduction of CSF-cN response. However, as RF loss is the only defect observed in the *scospondin*^*icm15*^ mutant (the dimensions of the CC and cilia motility are not altered in this mutant), we formulate the parsimonious assumption that the Reissner fiber is the important factor in explaining the loss of CSF-cN mechanosensory response.

### How can the RF and CSF-cNs interact to sense spinal curvature?

Recent evidence based on the labeling of SCO-spondin-GFP have revealed that the Reissner fiber *in vivo* is dynamic as well as straight as an arrow [47], which suggests that the fiber is under high tension *in vivo*. Given the dimensions of the central canal in living larvae (typically 10 µm by 9 µm) compared to the height of CSF-cN apical extensions (typically 3-5 µm) and the very thin diameter of the fiber (about 200 nm), it is conceivable that at rest, the thin RF could sit in the center of the central canal away from CSF-cN apical extension. In contrast, when muscles contract on one side (left, right or ventral), the Reissner fiber under tension could get closer to CSF-cNs during bending, which would lead to their selective asymmetrical recruitment (see Graphical Abstract).

The nature of the functional interaction between the Reissner fiber and CSF-cN apical extension could rely on a transient or stable physical contact, which would amplify the mechanical force applied on the apical extension of CSF-cNs in an asymmetrical manner during bending. Alternatively, from the fluid dynamics point of view, it is also highly conceivable that RF and CSF-cN apical extension *functionally* interact without the need for *physical* contacts. Indeed, the Reissner fiber could increase the CSF flow gradient perceived by CSF-cN apical extension. We previously showed that CSF flow in the central canal is maximal close to the center and null along the central canal walls [43]. Remarkably, CSF flow has to be null as well on the surface of the fiber itself. Therefore, CSF-cN apical extension pointing towards the center of the lumen sits in a region of high CSF flow gradient precisely at the boundary between the high flow and the null flow point imposed by the fiber. This effect could be amplified by that CSF flow is largely increased by muscle contractions along the tail as reported in the brain ventricles [48] and in the central canal [43].

Due to the fixation artifact that we quantified here with classical immunostaining protocol, further investigations of the dynamic interactions of CSF-cN apical extensions together with labeled RF *in vivo* [43] occurring during spinal curvature will be necessary to distinguish between these hypotheses.

### Comparison of CSF-cNs with inner ear hair cells

Kolmer and Agduhr originally compared CSF-cNs to hair cells due to the morphological similarity of the apical extension of both cell types. A century later, can we comment on the similarity of mechanisms underlying their mechanosensory functions? Of course, CSF-cNs with their coral-like shaped microvilli lack the regular staircase organization of stereocilia. However, CSF-cNs bear a kinocilium [15,16,49], similarly to inner ear hair cells from fish and amphibians [50]. We know from hair cells in amphibians that the active oscillations of the hair bundle amplify mechanical stimuli, which contributes to sound detection [51]. Similarly, through active movements, CSF-cN kinocilium could contribute to the amplification of mechanosensory response. The specific role of CSF-cN kinocilium in mechanoreception will be the focus of future investigations based on tools for disrupting cilia only in CSF-cNs and not in other cell types.

### Relevance for development of body axis and spine

Sensory systems are critical to guide symmetrical growth and balanced activation of motor circuits, but they can be also relevant for morphogenesis. We previously showed that CSF-cNs modulate the spinal circuits controlling locomotion and active posture [16,25,29,31,36]. Recently, the RF and CSF-cNs have also been associated with the establishment of the body axis during embryogenesis [8] and of the spine morphogenesis in juvenile and adult zebrafish [9,36]. Multiple evidence suggest that CSF-cNs are relevant for spine morphogenesis: *pkd2l1* mutants deprived of CSF-cN sensory responses exhibit an increased curvature of the spine, reminiscent of kyphosis [36]. Furthermore, mutants for a receptor of the urotensin related peptides, which are solely produced by CSF-cNs in the spinal cord, exhibit a torsion of the spine, reminiscent of adolescent idiopathic scoliosis [9]. Finally, a recent report indicated that hypomorphic mutations in the *scospondin* gene induce 3D deformation of the spine [47]. Altogether, recent studies indicate that sensory neurons contacting the cerebrospinal fluid together with the Reissner fiber may contribute to the generation and maintenance of the shape of the spine. Future studies will investigate whether the functional coupling of CSF-cNs with the Reissner fiber that we have demonstrated here is relevant for sensing and adjusting morphogenesis of the spine.

## Acknowledgments

We thank Monica Dicu and Antoine Arneau for fish care. We thank Céline Revenu for sharing the *Tg(cdh2:cdh2-GFP,crybb1:ECFP)* transgenic line; Remi Leborgne from the ImagoSeine facility (Jacques Monod Institute Paris, France), and the France BioImaging infrastructure supported by the French National Research Agency (ANR-10-INSB-04, " Investmentsfit the future”) for precious help in serial block face scanning electron microscope imaging. We thank for critical feedback all members of the Wyart lab (https://wyartlab.org/). Part of this work was carried out in the Phenoparc and ICM.Quant core facilities of ICM. This work was supported by an ERC Starting Grants “Optoloco” #311673 and, New York Stem Cell Foundation (NYSCF) Robertson Award 2016 Grant #NYSCF-R-NI39, the HFSP Program Grants #RGP0063/2014 and #RGP0063/2018, the Fondation Schlumberger pour l’Education et la Recherche (FSER/2017) for C.W. The research leading to these results has received funding from the program “Investissements d’avenir” ANR-10-IAIHU-06 (Big Brain Theory ICM Program) and ANR-11-INBS-0011 – NeurATRIS: Translational Research Infrastructure for Biotherapies in Neurosciences.

## Author Contribution

A.O.D.I. performed all calcium imaging and electrophysiology experiments and analysis, A.O.D.I. and Y.C.-B. performed immunohistochemistry and confocal microscopy imaging, A.B. and D.L. performed E.M., O.T. provided guidance on automated analysis for calcium imaging, J.R. and F.K. helped with genotyping, P.B. provided help for 3D reconstruction, F.-X.L. performed statistical analysis, P.-L.B. provided feedback on morphological analysis, C.W. conceived and supervised the project. A.O.D.I. and C.W. wrote the article with inputs from Y.C.-B., P.-L.B. and all authors.

## STAR Methods

### EXPERIMENTAL MODEL

All procedures were performed on 3 days post fertilization (dpf) zebrafish larvae in accordance with the European Communities Council Directive (2010/63/EU) and French law (87/848) and approved by the Institut du Cerveau et de la Moelle épinière (ICM). All experiments were performed on *Danio rerio* embryos of AB, Tüpfel long fin (TL) and nacre background. Animals were raised at 28.5°C under a 14 / 10 light / dark cycle until the start of the experiment.

**Table 1.**
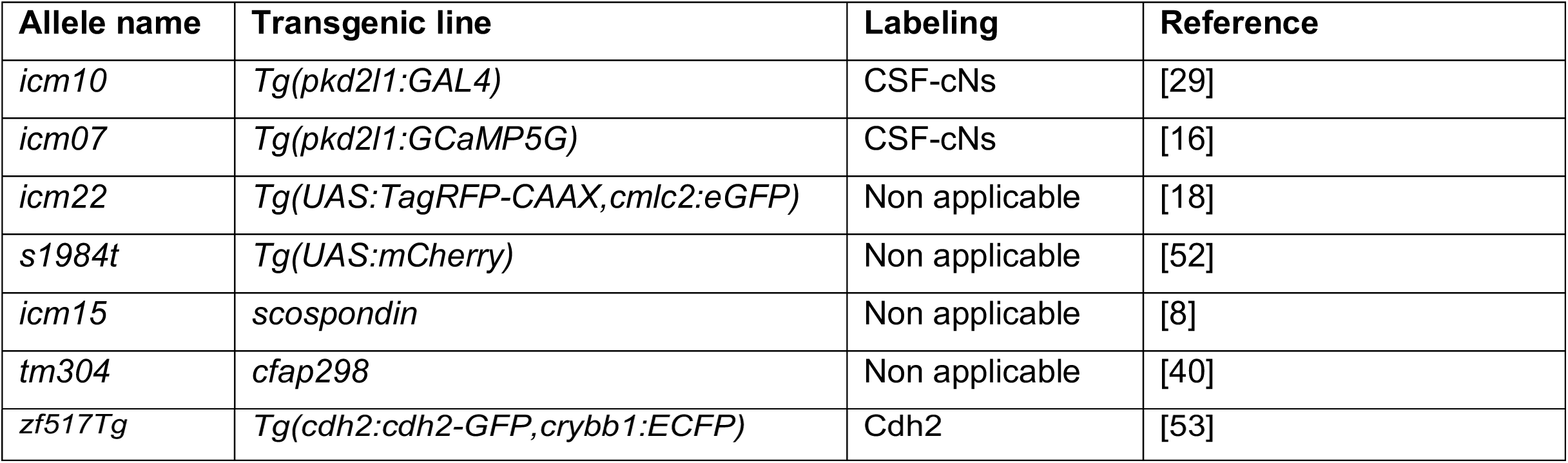
Mutant and transgenic lines used in our study.

### METHOD DETAILS

### Immunohistochemistry

3 dpf larva were euthanized in 0.2% Tricaine (MS-222, Sigma-Aldrich, Saint Louis, Memphis, USA) in system water and fixed 2 hours in phosphate-buffered saline (PBS) containing 4% paraformaldehyde (PFA) and 3% sucrose at 4°C. After three washes in PBS, skin from the rostral trunk was partially removed and the yolk was removed. Samples were incubated overnight in a blocking buffer containing 0.5% Triton, 1% DMSO, 10% normal goat serum (NGS) in PBS. Primary antibodies were incubated 48 hours at 4°C in a blocking buffer containing 0.5% Triton, 1% DMSO and 1% NGS. All secondary antibodies were from Molecular Probes© (Invitrogen, Life Technologies, Carlsbad, California, USA) unless specified and used at 1:500 in blocking buffer, and incubated 2.5 hours at room temperature. The following primary antibodies were used for *in toto* immunohistochemistry: rabbit anti-Reissner fiber (1:200, polyclonal, custom-made) [8,54], rabbit anti-Pkd2l1 (1:200, custom-made, Sternberg et al., 2018), mouse anti-tagRFP (1:500, MA515257, Thermo Fischer Scientific, Waltham, Massachusetts, USA), mouse anti-ZO-1 (1:200, 339100, Invitrogen) and chicken anti-GFP (1:500, ab13970, Abcam, Cambridge, England). The following secondary antibodies were used at 1:500: Alexa Fluor-568 goat anti-rabbit IgG A11036, Alexa Fluor-488 donkey anti-rabbit A21206, Alexa Fluor-647 goat anti-rabbit IgG A21244, Alexa Fluor-555 goat anti-mouse IgG1 A21127, Alexa Fluor-568 goat anti-mouse A11004, Alexa Fluor-488 goat anti chicken IgG A11039 (Thermo Fischer Scientific, Waltham, Massachusetts, USA). Zebrafish larvae were mounted dorsally (for imaging in coronal plane) or laterally (for imaging in sagittal plane) in Vectashield® Antifade Mounting Medium (Vector Laboratories, Inc., Burlingame, California, USA) and imaged on an Olympus FV-1000 confocal microscope equipped with a 40X NA = 1.3 oil immersion objective (0.5 µm optical section and 0.35 µm optical section for images used to count contact with the RF (see Figure 5). Images were then processed using Fiji [55].

### Analysis of the distribution of the apical extensions of cerebrospinal fluid contacting neurons in close vicinity with the Reissner fiber

To image the Reissner fiber together with cerebrospinal fluid contacting neurons membranes in *Tg(pkd2l1:GAL4;UAS:tagRFP-CAAX)*, zebrafish larvae were mounted dorsally or laterally in Vectashield Antifade Mounting Medium (Vector Laboratories, Inc., Burlingame, California, USA) and imaged on an SP8 X White Light Laser Leica inverted confocal microscope equipped with a 63X oil immersion objective (NA = 1.4). Dorsally mounted larvae allowed acquiring coronal planes ideally oriented to access the apical extensions of dorsolateral cerebrospinal fluid contacting neurons, while laterally mounted larvae allowed acquiring sagittal planes ideally oriented to access the apical extensions of ventral cerebrospinal fluid contacting neurons (see Figure 5). Z-stacks with a 250 nm step size (pixel size in the (x,y) plane: 144 nm) were acquired to estimate the proximity of the Reissner fiber with the apical extensions of cerebrospinal fluid contacting neurons in both cases. Four consecutive spinal cord regions were acquired along the rostro-caudal axis of the animals, respectively between segments 5 and 16. In order to correct for optical distortions taking place in the (x,y) and (z) planes, Z-stacks were first deconvolved for both the Reissner fiber fluorescence signals and cerebrospinal fluid contacting neurons membrane signals, with the Huygens Professional version 19.04 (Scientific Volume Imaging, The Netherlands, http://svi.nl), using the CMLE algorithm, with a signal-to-noise ratio between 15 and 20, and up to 40 iterations. 3D stacks were then processed using Fiji [55]. Apposed or colocalized immunofluorescence signals from CSF-cNs and the Reissner fiber at least in a single plane after deconvolution were considered in close vicinity. The percentage of apical extensions in close vicinity with the Reissner fiber was then estimated for each fish. 3D views of the Reissner fiber and CSF-cNs from coronal and sagittal views (Figure 5E) were obtained using the 3D Viewer plugin on Fiji [56], after applying a median filter prior to 3D-segmentation.

### Morphological Analysis

#### Analysis of the shape of the central canal in live and fixed 3 dpf larva

To measure the size of the central canal, we used 3 dpf zebrafish larvae where the apical junctions of the ependymal cells were visualized (with ZO-1 or Cadherin-2). For live imaging, we used the 3 dpf double transgenic *Tg(cdh2:cdh2-GFP; pkd2l1:GAL4;UAS:tagRFP-CAAX)* larva [53] in order to visualize the central canal and CSF-cNs *in vivo*. For images on fixed animals, we used antibodies against GFP (for cdh2-GFP), ZO-1 and tag-RFP. All images were acquired with an SP8 X White Light Laser Leica inverted confocal microscope equipped with a 40X water immersion objective (NA = 1).This image analysis was performed with Fiji. For the height of the canal, we acquired z-stacks of sagittal confocal sections spaced by 1 µM. A max z-projection of the slices encompassing the canal (typically 5 to 10 slices) was performed, and the height of the fluorescent signal what measured at 4 different fixed levels (spaced by 30 µM) and averaged. For the width of the canal, we acquired z-stacks of coronal confocal sections spaced by 1 µM. We observed frequently that the canal was not straight in this axis, preventing us from quantifying the width on z-projection. Instead, we quantified at 4 different fixed positions (spaced by 30 µM) the width of the fluorescent signal in the 5^th^ slice above the floor plate signal (hence 4 µM above the floor plate), and averaged it.

#### Measurement of the size of CSF-cN apical extension in vivo

To measure the size of the apical extension, we used 3 dpf *Tg(cdh2:cdh2-GFP; pkd2l1:GAL4;UAS:tagRFP-CAAX)* zebrafish larvae where CSF-cNs were visualized through RFP signal. All images were acquired with an SP8 X White Light Laser Leica inverted confocal microscope equipped with a 40X water immersion objective (NA = 1). To measure the height of the apical extension, we acquired z-stacks of sagittal confocal sections spaced by 1 µM. A max z-projection of the slices encompassing each CSF-cNs was performed and we drew polygons outlining their apical extension using the polygon tool in Fiji.sc as described previously. An ellipse was fitted to the polygon and the height of the apical extension was measured as the height of the axis of the ellipse perpendicular to the floor plate as described previously [17,18].

### Calcium imaging

All experiments were done on 3 dpf *Tg(pkd2l1:GCaMP5G); scospondin*^*icm15/icm15*^ and *Tg(pkd2l1:GCaMP5G); cfap298* ^*tm304/tm304*^ and their respective siblings used as controls (*i.e.* wild type and heterozygous mutants from the same clutch).

#### Active muscle contraction

Unparalyzed 3 dpf larvae were pinned on their side through the notochord with 0.025 mm tungsten pins and bathed in artificial cerebrospinal fluid solution (aCSF, concentrations in mM: 134 NaCl, 2.9 KCl, 1.2 MgCl_2_, 10 HEPES, 10 glucose and 2.1 CaCl_2_; 290 mOsm.kg^−1^, adjusted to pH 7.7–7.8 with NaOH). Active muscle contraction was induced by a 1 s-long pressure application of aCSF on the otic vesicle repeated 4-5 times with an inter trial interval of 1 minute. GCaMP5G fluorescence was excited by 490 nm illumination and monitored for 250 s at 4 Hz using an Examiner epifluorescence microscope (Zeiss, Göttingen, Germany) equipped with a 40 X NA = 1.0 water-immersion objective and EMCCD camera ImagEM X2 (Hamamatsu, Naka-ku, Japan). Images were acquired using Labview software (National Instruments, Austin, Texas, USA) for *cfap298*^*tm304*^ mutants and using Hiris software (R&D Vision, Nogent-sur-Marne, France) for *scospondin*^*icm15*^ larvae and reconstructed using Fiji.

#### Passive spine bending

3 dpf larvae were anesthetized in 0.02% Tricain (MS-222, SIGMA dorsally mounted in glass-bottom dishes (MatTek, Ashland, Massachusetts, USA) filled with 1.5% low-melting point agarose. Larvae were paralyzed by injecting 0.5 nl of 0.5 mM α-Bungarotoxin in the musculature (Tocris Bioscience, Bristol, UK, [57] and placed in aCSF. After embedding, roughly half of the larval tail was freed unilaterally to provide access to a blunt 50 μm diameter glass probe. Probe deflections were driven with a mechanotransducer device controlled through LabView software as done previously [16,17]. Calcium imaging was performed on a two-photon laser scanning microscope (2p-*vivo*, Intelligent Imaging Innovations, Inc., Denver, USA) using a 20X NA = 1.0 objective. Lateral bending of the tail was induced by probe deflection and repeated 3 times every 14 to 17.5 s as done previously [16,17].

#### Calcium imaging analysis

Slow translational drifts of the image due to spine movement were corrected using image registration by taking as a reference image a max Z-projection of 3 consecutive images chosen when CSF-cNs are back to their position and bright after a muscle contraction. The regions of interest (ROI) corresponding to each individual cell were drawn manually on the reference image. We identified CSF-cN calcium transients in response to spinal curvature by using either the motion artifact itself or a 200 ms-long flash of green light performed 16 s before the stimulus. The amplitude of the first CSF-cN calcium transients in response to passive and active spinal concave curvature were determined relative to baseline preceding the motion artifact with custom scripts written in MATLAB (MathWorks, Natick, Massachusetts, USA). For each contraction and each ROI, ΔF / F was estimated as (F_GCaMP_ - F_0-GCaMP_) / F_0-GCaMP_ with F_GCaMP_ is the fluorescence signal averaged over four time points (*i.e.* 1 s) around the peak after the first contraction and F_0-GCaMP_ is the baseline fluorescence average over 3 time points (*i.e.* 0.75 s) before each motion artifact. For each contraction, a new baseline was therefore estimated to prevent errors due to photobleaching or drifting during the recording. The percentage of responding cell was set as ΔF / F > 1.96 STD of the minimum value during the motion artefact.

### In vivo patch-clamp recording

Whole-cell recordings were performed in aCSF on 3 dpf *Tg(pkd2l1:Gal4; UAS:mCherry)* carrying either the *scospondin*^*icm15*^ or *cfap298*^*tm304*^ mutation and their respective control siblings. Larva were pinned through the notochord with 0.025mm diameter tungsten pins. Skin and muscle from two to three segments around segment 10 were dissected out using a glass suction pipette. A MultiClamp 700B amplifier, a Digidata series 1440 A Digitizer, and pClamp 10.3 software (Axon Instruments, Molecular Devices, San Jose, California, USA) were used for acquisition. Raw signals were acquired at 50 kHz and low-pass filtered at 10 kHz. Patch pipettes (1B150F-4, WPI) with a tip resistance of 5–8MΩ were filled with internal solution (concentrations in mM: K-gluconate 115, KCl 15, MgCl2 2, Mg-ATP 4, HEPES-free acid 10, EGTA 5 or 10, 290 mOsm/L, pH adjusted to 7.2 with KOH with Alexa 488 at 40 μM final concentration). Holding potential was −85 mV, away from the calculated chloride reversal potential (E_Cl_ = - 51 mV). Analysis of electrophysiological data was performed offline using Clampex 10 software (Molecular Devices, San Jose, California, USA). Single channel events were identified using single-channel search in Clampfit (Molecular Devices, San Jose, California, USA), with a first level set at −15 pA from the baseline (level 0). Only events lasting longer than 1.2 ms were included for analysis. A 20 s window was used to identify channel events from a gap-free voltage-clamp recording from the first 1 to 3 min of recording. Passive properties were determined, in voltage-clamp mode at −85 mV, from the cell current response to a 10 mV hyperpolarization step (V step). Membrane resistance (R_m_) was estimated from the amplitude of the sustained current at the end of the 100 ms voltage step (R_m_ = V step / Im). Membrane capacitance (C_m_) was estimated as the ratio between the cell decay time constant (t), obtained from the exponential fit of the current decay an R_S_ (C_m_ ∼ t / R_S_). Action potential discharge was monitored in current-clamp mode in response to successive depolarizing current steps of 100 ms from – 2 pA to + 28 pA steps with a 2 pA increment after a fixed prepulse with - 10 pA for 20 ms while holding the cell membrane potential at - 50 mV.

### Electron microscopy

All the products used for electron microscopy were obtained from Electron Microscopy Science (EMS, distributor Euromedex, Souffleweyesheim, France).

#### Transmission electron microscopy

Samples were fixed in 0.5% glutaraldehyde 4% PFA in PBS, pH 7.4 for 2 hours at 4 °C. Some samples were treated with 1% trichloroacetic acid (TCA) within the fixative solution in order to better visualize preserve the Reissner fiber (as shown in Figure 6A1, 6A2, 6A3). Following three rinses with Na-cacodylate buffer 0.1 M pH = 7.4, sections were post-fixed with 1 % osmium tetroxide in the same buffer for 1 hour. Samples were dehydrated in a graded series of ethanol solutions (75, 80, 90 and 100 %, 5min each). Final dehydration was performed twice in 100 % acetone for 20 min. Infiltration with epoxy resin (Epon 812) was performed in 2 steps: overnight at + 4°C in a 1:1 mixture of resin and acetone in an airtight container and then, 2 hours at room temperature (RT) in pure resin. Finally, samples were placed in molds with fresh resin and cured at 56°C for 48 hours in a dry oven. Samples were sagittally cut in 0.5 µm semi-thin sections with an ultramicrotome EM UC7 (Leica, Wetzlar, Germany). Sections were stained with 1% toluidine in borax buffer 0.1 M. Then ultra-thin sections (∼ 70nm thick) were cut and collected on copper grid. Sections were then contrasted with Reynolds lead citrate for 7min[58]. Observations were made with a HT 7700 electron microscope operating at 70kV (Hitachi, Ltd, Tokyo, Japan). Electron micrographs were taken with an integrated AMT XR41-B camera (2048 x 2048 pixels, Advanced Microscopy Techniques Corp., Woburn, Massachusetts, USA).

#### Serial block face scanning electron microscope imaging (SBF-SEM)

Samples (as shown in Figure 6C and Supplemental Video 3) were fixed in 0.5% glutaraldehyde 4% PFA in PBS pH 7.4 for 2 hours at 4 °C. Following three rinses with Na-cacodylate buffer 0.1M pH = 7.4, sections were post-fixed for 30 minutes with 0.1% tannic acid in caco buffer as a mordant. After three rinses in caco buffer, samples were stained in freshly prepared (1% OsO_4_; 0,15% K_4_Fe(CN)_6_) solution for 1 hour. Samples underwent multiple incubation steps to increase the contrast: 20 min in 0.01% thiocarbohydrazide (TCH) at 60°C, 30 minutes in 1% OsO_4_, 60 min at 4°C in 1% aqueous uranyl acetate and 30 min in a 0.66% lead nitrate in 30mM aspartic acid solution, pH = 5.5 at 60°C (Walton, 1979). Samples were dehydrated in graded ethanol at room temperature with a final dehydration in 100% acetone. Samples were then embedded in 50% resin / 50% acetone overnight at +4°C and dry at 60°C for 48 hours. Samples were sagittally sectioned in 7 nm-thick sections every 40 nm and imaged with a SBF-SEM. Sectioning and scanning were performed with a TeneoVS electron microscope (FEI Company, Hillsboro, Oregon, USA) operating at 2kV-100pA-low vacuum (40Pa)-dwell time 1µs. Subsequently, 3D reconstruction was made using the Imaris software (Oxford instruments, Zurich, Switzerland).

### STATISTICAL ANALYSIS

All values are mean ± standard error of the mean (S.E.M.) and represented as a box plot where the central mark on indicates the median, the bottom and top edges of the box indicate the 25th and 75th percentiles. The whiskers extend to the most extreme data points that are not considered outliers, outliers are identified with a “+” symbol.

### CSF-cN Patch-clamp recording and Morphological study

Statistical significance was determined using Two-sample Kolmogorov-Smirnov test (kstest2, MATLAB, MathWorks, Natick, Massachusetts, USA). A value of p ≤ 0.05 was considered significant.

### Calcium imaging

Stimulus artifact have been digitally removed in all figures.. The data were analyzed with the repeated measure design. Values obtained from the response to active stimulation were analyzed using linear mixed-effects models (LMMs) with condition (control *vs.* mutant) and domain (dorsolateral *vs.* ventral CSF-cNs) as fixed effects and each independent fish (nested within clutch) as a random effect to account for the repeated measurements. Significance for the main effects of condition, domain and their interaction were then evaluated using ANOVA Type II Wald chi-square tests. The same analysis was conducted with the values obtained from response to passive stimulation, but for the condition factor only. All statistical analyses were conducted using R version 3.5.2 [59] and plots were generated with the ggplot2 package. All LMMs were fitted using the function lmer in the lme4 package. ANOVA Type II Wald chi-square tests were performed using the function anova in the car package. Post hoc Tukey’s comparisons of the conditions within domains were made and plotted using the estimated marginal means from the emmeans package. To improve normality and homoscedasticity of residuals in the LMMs, response data were square root transformed on absolute values and then returned to their original sign prior to analysis. The level of statistical significance was set at *p* < 0.05 for all tests.

### DATA AND CODE AVAILABILITIES

Further information and requests for resources and codes should be directed to and will be fulfilled by the Lead Contact, Claire Wyart.

### KEY RESOURCES TABLE

#### Key Resources table

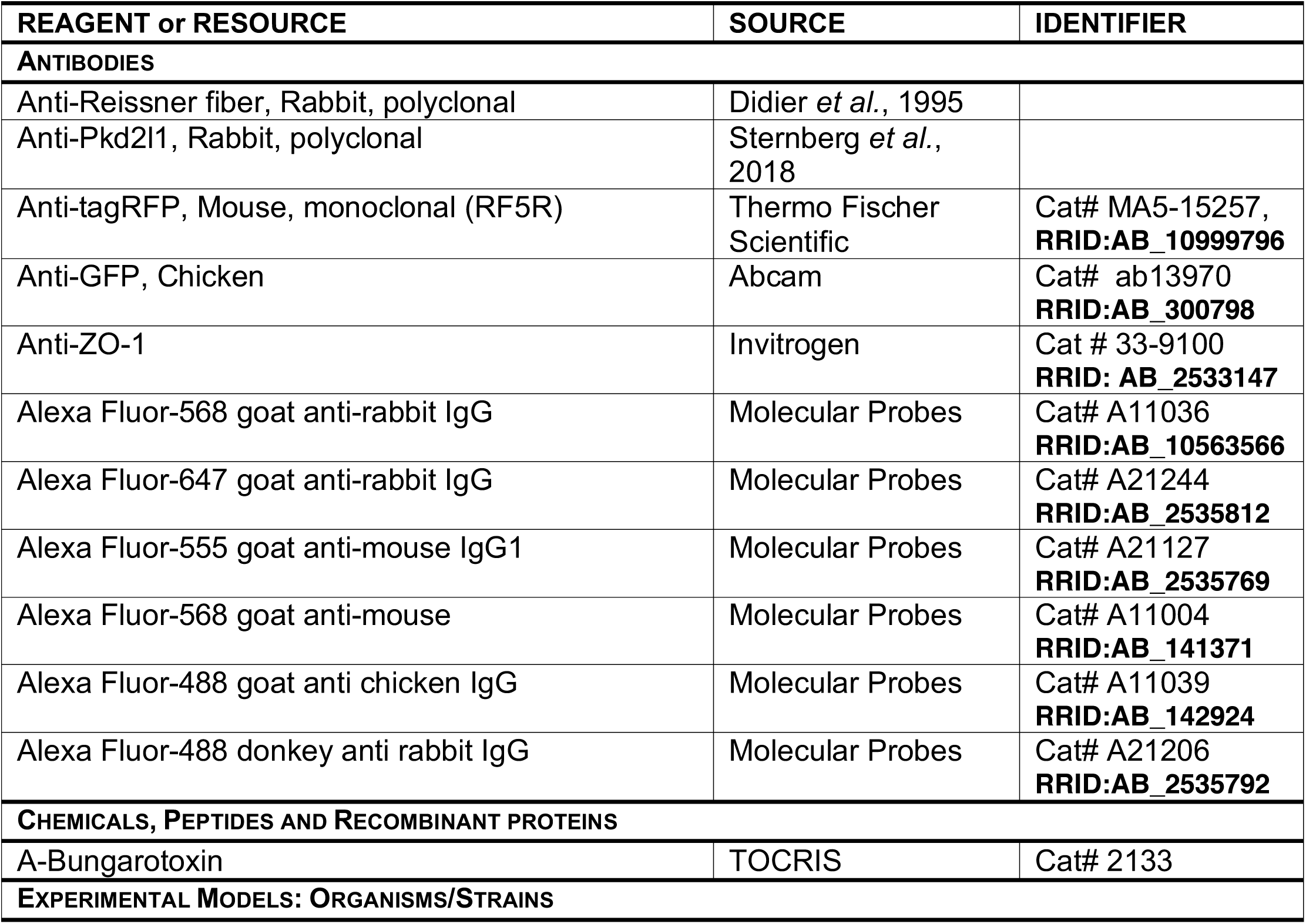
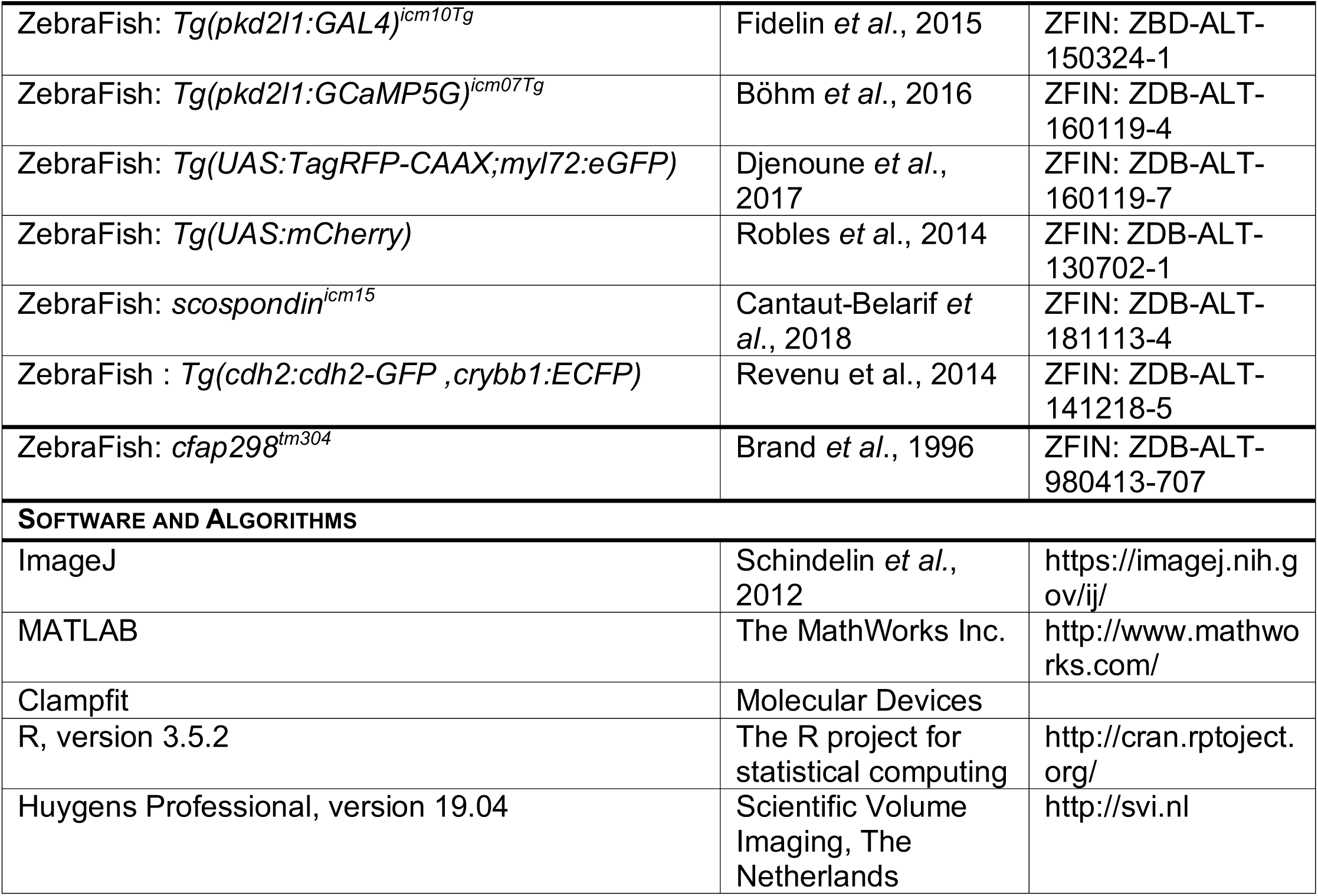

## Supplemental Movies

**Supplemental Movie S1. Large calcium transients occur in CSF-cNs upon spinal curvature. Related to Figure 1.** Large calcium transients in dorsolateral and ventral CSF-cNs were induced by active muscle contractions after pressure-application of aCSF on the otic vesicle. Lateral view of the spinal cord in a 3 dpf *Tg(pkd2l1:GCaMP5G)* larva in which calcium imaging (raw data) was acquired at 4 Hz and is displayed at 30 Hz. Motion artifacts where cells are moving during contractions are excluded for subsequent analysis of ΔF/F.

**Supplemental Movie S2. CSF-cN mechanosensitivity is disrupted in the *cfap* mutant with defective ciliary motility. Related to Figure 1.** Calcium transients in CSF-cNs induced by active muscle contractions after pressure-application of aCSF on the otic vesicle are massively reduced in *Tg(pkd2l1:GCaMP5G); cfap* ^*tm304/tm304*^ larvae. Lateral view of the spinal cord in which calcium imaging (raw data) was acquired at 4 Hz and is displayed here at 30 Hz. Motion artifacts where cells are moving during contractions are excluded for subsequent analysis of ΔF/F.

**Supplemental Movie S3. The absence of the RF disrupts CSF-cN responses to spinal curvature. Related to Figure 3.** Calcium transients in CSF-cNs upon active muscle contractions after pressure-application of aCSF on the otic vesicle are massively reduced in Tg(pkd2l1:GCaMP5G); *scospondin* ^*icm15/icm15*^ larvae. Lateral view of the spinal cord in which calcium imaging (raw data) was acquired at 4 Hz and is displayed here at 30 Hz. Motion artifacts where cells are moving during contractions are excluded for subsequent analysis of ΔF/F.

**Supplemental Movie S4. The RF is in close proximity with cilia and CSF-cN apical extension in the central canal. Related to Figure 6.** SBF-SEM imaging (60 sections of 7nm-thickness and with 40 nm-Z step) is displayed at 15 Hz and show the organization of RF in the lumen of the central canal.

